# Neuronal Mechanisms of Strategic Cooperation

**DOI:** 10.1101/500850

**Authors:** Wei Song Ong, Seth Madlon-Kay, Michael L. Platt

## Abstract

Here we demonstrate that during strategic gameplay monkeys behave as if they reason recursively about other individuals’ beliefs and desires in order to predict their choices and to guide their own actions, especially the decision to cooperate. Neurons in mid superior temporal sulcus (mSTS), the putative homolog of the human temporo-parietal junction (TPJ), signal abstract non-perceptual social information, including payoffs, intentions, and outcomes, and further distinguish between social and nonsocial agents while monkeys play the game. We demonstrate for the first time that a subpopulation of these neurons selectively signals cooperatively obtained rewards. Neurons in the anterior cingulate gyrus (ACCg), an area implicated in vicarious reinforcement and empathy, do not distinguish agency and as a population carry less information about strategic variables. These findings suggest the capacity to mentalize has deep roots in the strategic social behavior of primates, and endorse mSTS as the evolutionary wellspring of these functions.

---

Both emotional and cognitive mechanisms shape the decisions people make when they interact with others ^1,2^. Specifically, vicarious feelings of reward or pain experienced by another, often termed empathy, can provoke prosocial actions ^3^. Strategic reasoning about the beliefs, desires, and goals of another individual, a process referred to as mentalizing or theory of mind, guides the decision to cooperate with or betray a partner ^4,5^. These two processes interact as well; manipulations that increase empathy enhance cooperation ^6^. Two separate but interacting brain systems appear to support empathy and mentalizing during social decisions ^7^. In humans, empathy and vicarious experience evoke hemodynamic activity in anterior cingulate gyrus (ACCg), anterior insula, and amygdala, and neurons in primate ACCg and amygdala signal vicarious rewards delivered to other monkeys ^8,9^. By contrast, thinking about the beliefs, desires, or goals of others evokes hemodynamic activity in the dorsomedial prefrontal cortex and temporo-parietal junction (TPJ) in humans ^7,10–12^. The neuronal mechanisms underlying such mentalizing-related brain activity, however, remain poorly understood in part due to the difficulty of eliciting recursive social reasoning in primates or other animals in which neuronal activity can be studied directly (but see Haroush^13^) as well as the lack of neurophysiological or histological evidence for a TPJ homolog in nonhuman primates ^14,15^.

To address this gap, we trained monkeys to play a version of the classic “chicken” game from behavioral economics ^16^. We also recorded spiking activity of 448 neurons in the middle superior temporal sulcus (mSTS), a brain area known to encode perceptual social information like faces ^17,18^ and recently proposed as the primate homolog of TPJ based on MRI-based functional connectivity ^15^. For comparison, we recorded spiking activity of 528 neurons in ACCg, an area strongly linked to vicarious reward and empathy ^8,19,20^

Our variant of the chicken game allowed players to coordinate in pursuit of a cooperative reward, as well as pursue individual rewards at the expense of the other player. The sizes of the cooperative and individual payoffs changed on each trial, encouraging animals to dynamically switch between competing and cooperating. In each play session, a monkey played against either another monkey, a computer, or a computer with a decoy monkey present. In the live and decoy conditions, two monkeys faced each other over a screen that was placed horizontally between them and parallel to the ground (Figure 1a). They used joysticks to interact with the game and eye position was recorded at 1000 Hz (Eyelink). Two colored rings framing moving dots (hereafter ‘cars’) and 6 arrays of tokens were presented on the screen. Token arrays indicated the amount of juice reward available for going straight or deviating alone for each player. If one player went straight and the other deviated, each would receive juice proportional to the tokens acquired. If both players went straight, the cars “crashed” into each other, and no reward was delivered. If both monkeys chose to deviate they received the associated rewards plus bonus tokens released by pushing the cooperation bar (Figure 1c & 1f). Payoffs varied randomly from trial to trial. The white dots within the car flowed in the direction in which the joystick was currently held, providing an intention cue to the opponent that could either be clear (100% correlated dots) or ambiguous (0% correlated dots). When a player held the joystick in one direction for 0.5 secs, the dots changed color and the player’s choice was locked (see supplementary task video).

**Figure 1:**
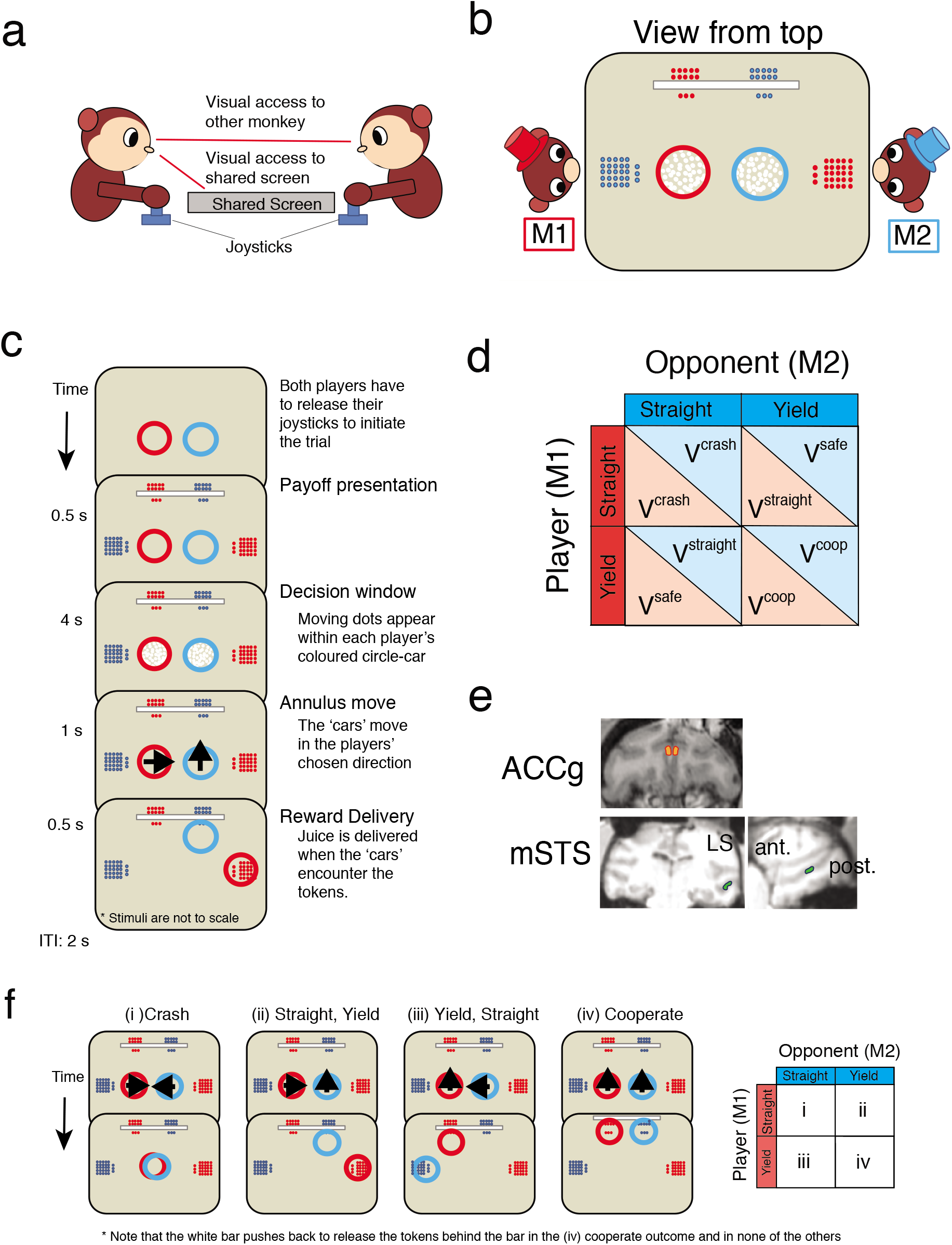
“Chicken” game with cooperation option. A. Two players (M1 & M2) sat opposite each other over a shared horizontal screen.
B. M1controlled red annulus (hereafter “car”) with joystick; M2 controlled blue annulus.
C. Task sequence.
D. Payoff matrix. Red player (M1) occupies row and blue player (M2) occupies column.
E. Recording sites in ACCg (orange) and mSTS (green).
F. Example outcomes for different actions. Each player could choose straight or yield.

Overall, monkeys made choices that aligned with the mixed strategy Nash equilibrium prediction based on available payoffs (Figure 2c) when the opponent’s intentions were clearly signaled by the moving dots in the cars. When intentions were ambiguous, only 69% of trial outcomes followed the mixed strategy Nash equilibrium (figure 2d). The fact that monkeys largely avoided crashing even in the absence of explicit intention signals suggests they relied on other information to guide their choices. We explored the possibility that monkeys used visual cues available from the other monkey and also tested the idea that monkeys formed a model of their opponent to guide their choices.

**Figure 2:**
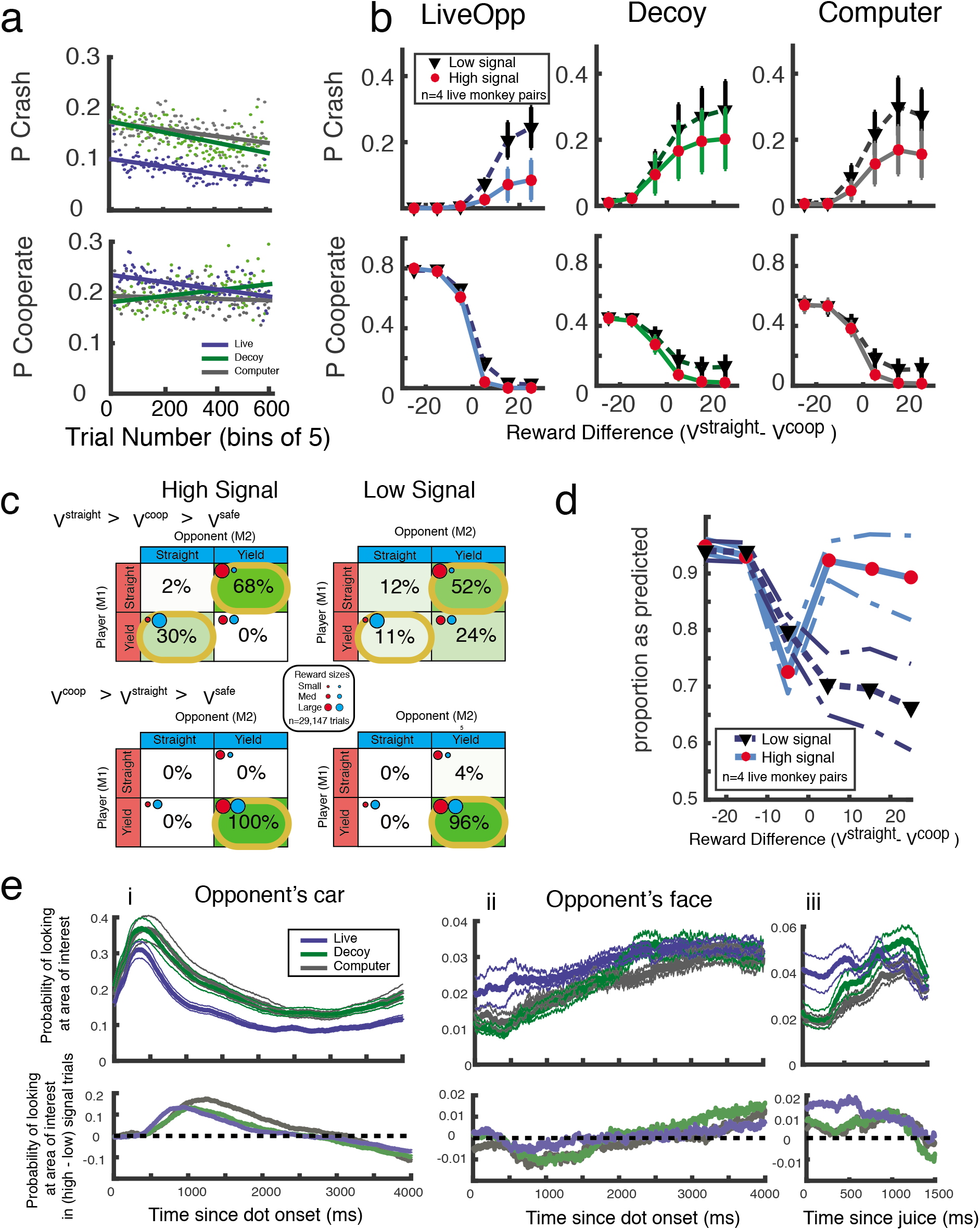
A &B) Monkeys understand the task and discriminate agency. Probability of crashing (top) or cooperating (bottom) over time and (B) by payoff difference for straight (Vstraight) and cooperate (Vcoop). (+-SEM). C) Monkeys look at most informative stimuli, calibrated by agency. (i) Top: Probability M1 looked towards opponent’s car synchronized to moving dots onset. SEM calculated between sessions. Bottom: Difference between high and low signal trials. (ii) Same as (i) but for opponent’s face. (iii) Same as (ii) but synchronized to juice delivery. D) Outcomes for a pair of live opponents across all sessions segregated by payout and signal strength. Yellow borders indicate Nash equilibria. E) Proportion of trials resulting in Nash outcomes indicated in 2D (+-SEM)

After a brief fixation period to start each trial, gaze was unconstrained. Monkeys spent most of the token onset period (500ms) looking at the tokens in front and to the side of the screen regardless of agency condition (supplementary figure 1a &b). Monkeys spent at least 1/3 of the first 500ms of the choice period looking at the opponent’s car (Figure 2ei, top panel). However, monkeys spent less time looking at the opponent’s car in the live condition than either the decoy or computer condition. This difference was offset by spending more time looking at the face of a live opponent, although monkeys also spent more time looking at a decoy’s face toward the end of the choice period. Thus, monkeys looked at key sources of information during the trial and their gaze distinguished agency conditions that were perceptually similar (decoy vs. live). Gaze also reflected whether intentions were signaled within the cars. Monkeys preferentially looked at the opponent’s car when signal strength was high compared with when it was low (Figure 2ei, lower panel). During reward delivery, monkeys were much more likely to look at a live opponent drinking earned juice than towards a decoy, and were more likely to look at any monkey than the dripping juice tube present in the computer condition (figure 2eiii). Finally, monkeys were much less likely to look at a monkey with whom he had cooperated to acquire bonus reward (supplementary figure 1c). Thus, monkeys adaptively sampled visual information about payoffs, the intentions of the opponent, and perceptual social cues, and this process betrayed a sense of the agency of the opponent.

These data invite the hypothesis that monkeys sampled multiple sources of information to compute and update a model of the opponent, which they used to guide their own choices. We explored this hypothesis by comparing their behavior to a series of decision models of increasing cognitive sophistication. Each model assumes that monkeys calculated the expected value of each option based on a prediction of his opponent’s actions. If his opponent was likely to go straight, he should yield to secure the small but safe reward instead of risking a crash. In the least sophisticated model, the monkey believes his opponent chooses with some fixed probability that is not influenced by the payoffs. In the more sophisticated models, the monkeys either realize their opponents also try to maximize their own payoffs and accordingly choose differently when payoffs are different, learn adaptively about their opponent’s strategies based on experience, or both. The learning models update beliefs about the opponent’s strategies using a strategic prediction error (SPE), the difference between the opponent’s predicted strategy and his actual choice. The best-fitting model was the most sophisticated, including both representation of the opponent’s utility and SPE-driven learning (mean decrease in AIC = 774) ^21^ (Figure 3b). For comparison, players’ behaviors did not follow tit-for-tat ^22^ or win-stay-lose-shift ^23^ strategies (supplementary figure 2b), and we found no evidence for simple reinforcement-learning.

**Figure 3:**
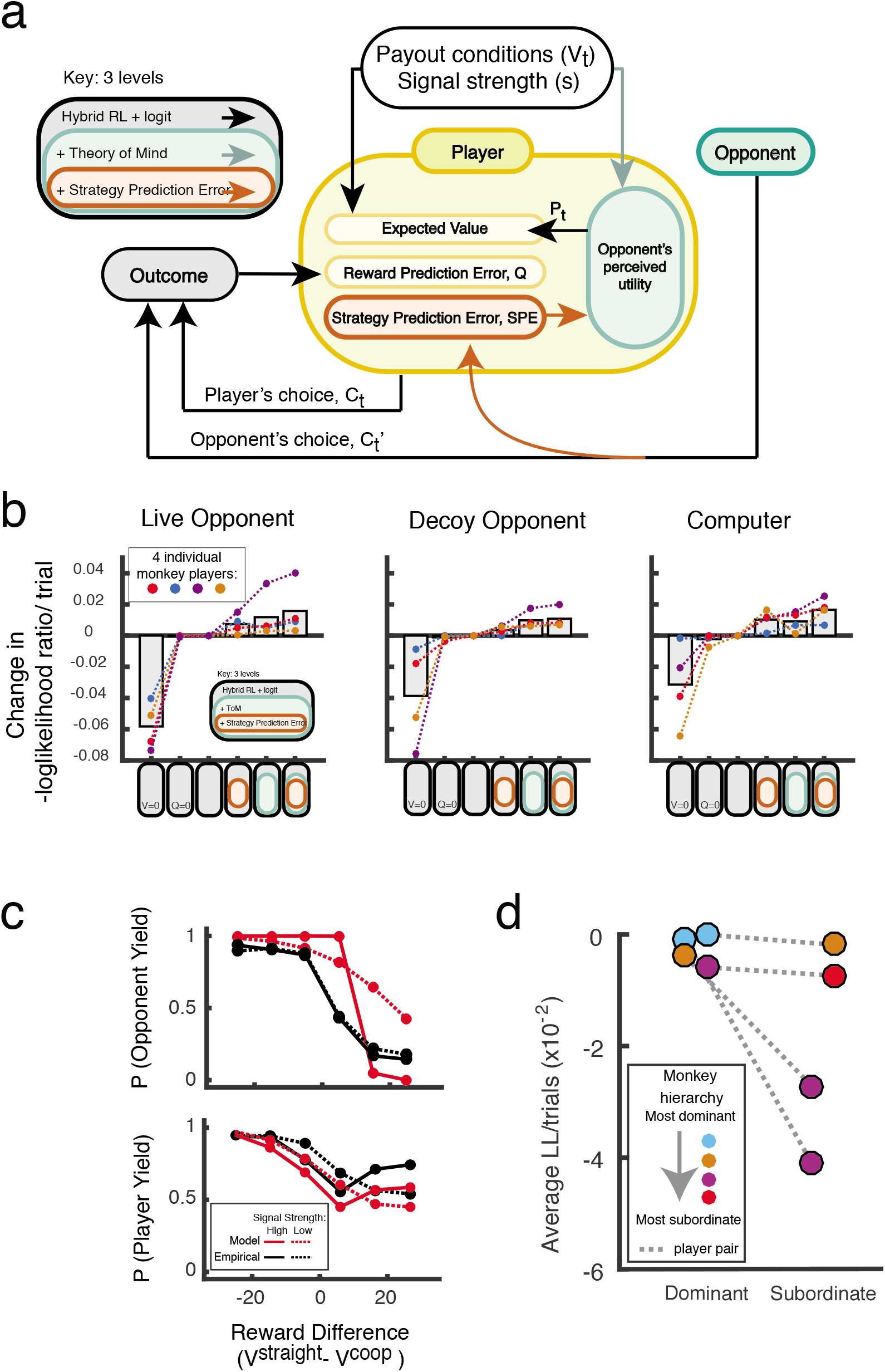
Model of player behavior and AIC values for each model. A. Model of events and variables that contribute to player’s internal calculations when making choices. For details see Methods.
B. Model comparisons for live, decoy, and computer conditions. Average log-likelihood ratio (LLR) change compared to basic model (grey bars) with exclusion of payout conditions (Vt), reward prediction error (Q) and inclusion of intentionality (k1) and strategic prediction errors (SPE).
C. ToM model fits and observed behavior for one player pair.
D. Model improvements vary with relative dominance status of players in a dyad.

Our modeling exercise suggests monkeys behave as if they reason recursively about other individuals’ beliefs, motivations, and strategy in order to predict their choices and to guide their own actions. The depth of this recursion depended on monkey identity (figure 3D). When intentionality was assigned to the opponent within the model, improvement in fit was greater for subordinate monkeys than for dominant monkeys. These findings suggest subordinate monkeys were more sensitive to the intentions of dominant monkeys in the game, consistent with prior reports that subordinate monkeys pay more attention to dominant monkeys, who themselves attend selectively to other dominant monkeys ^24,25^. Furthermore, the same mid-ranking monkey played against different opponents (brown and purple in figure 3d), and their strategies were more consistent with relative dominance than individual identity.

Our behavioral and eye-tracking data demonstrate monkeys are exquisitely sensitive to payoffs for self and their opponent, information about intentions, and reward outcomes, as well social information reflecting identity, social dominance, and perhaps gaze direction. Monkeys use this information to compute a model of their opponent, including how likely he is to behave cooperatively.

We next queried the role of neurons in ACCg, a brain area associated with vicarious reinforcement and empathy, and mSTS, a brain area linked to perceptual social processes and recently proposed as the primate homolog of human TPJ, in the computational processes underlying behavior in our task. We found that firing rates of neurons in both areas were sensitive to payoffs early in the trial, the presence of intention signals within the cars, and the amount of reward received after both monkeys made their choices. The strength and abundance of these signals varied between brain areas, across agency conditions, and as a function of time during each trial (Figures 4c, Supplementary Figure 3b).

**Figure 4:**
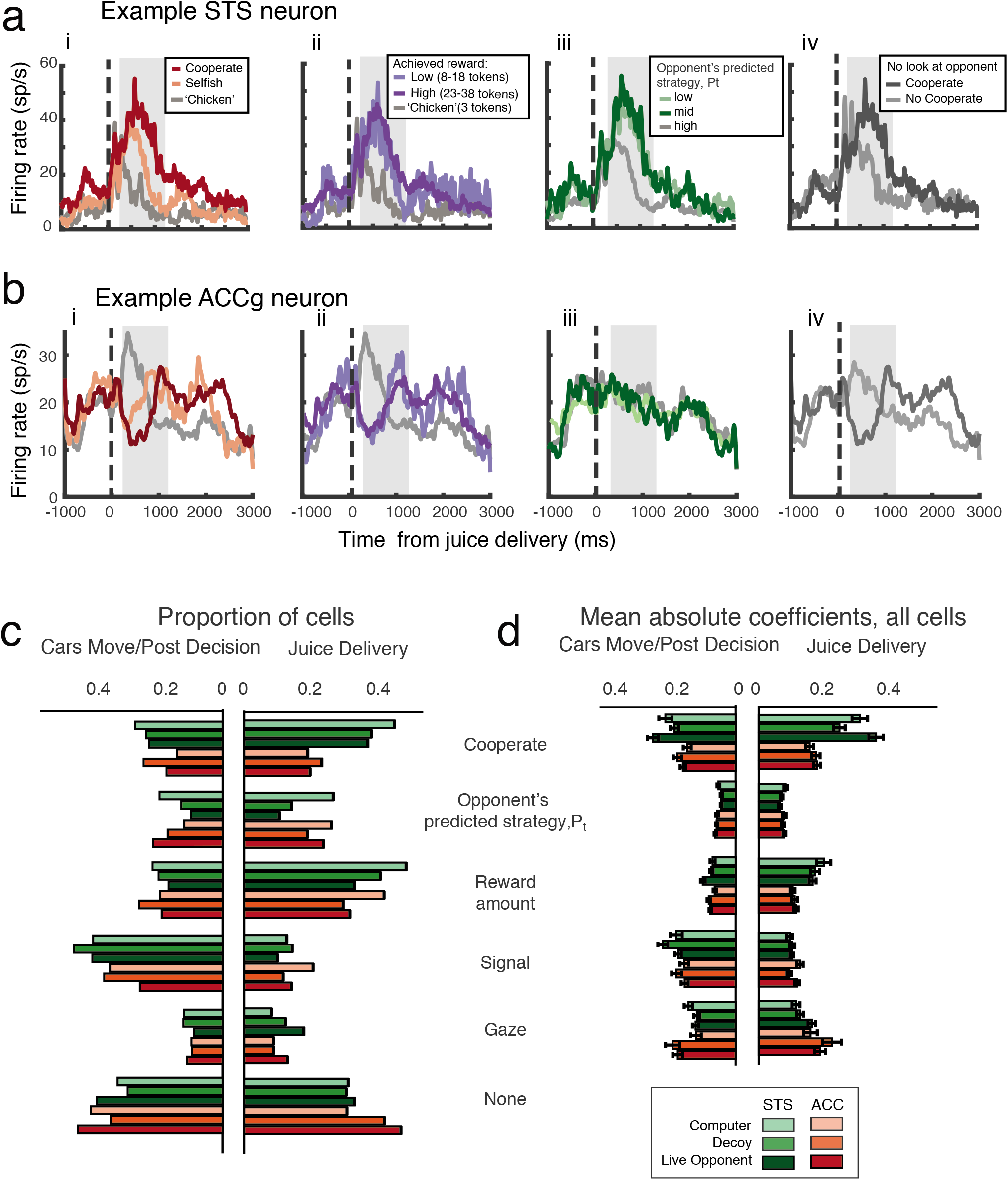
mSTS and ACCg neurons signal abstract, non-perceptual strategic information. A. PSTH for an mSTS neuron during play with live opponent. (i) Response to cooperative, non-cooperative and ‘chicken’ rewards. (ii) Responses to high, low and chicken rewards. (iii) Responses to high, mid, and low strategy prediction (Pt). (iv) Responses to cooperative vs non-cooperative rewards with no social gaze.
B. PSTH for an ACCg neuron during play with a live opponent. Conventions as in Figure 4A.
C. Percentage of cells in ACCg and mSTS in 3 agency conditions showing significant modulation by signal strength, reward size, gaze, Pt, and cooperation (0-500ms after car movement and 250-1250ms after reward. Also see supplementary table 1.
D. Mean absolute model coefficients for all neurons.

We used linear models (LMs) to quantify neuronal sensitivity to payoffs, intention signals, reward outcomes, cooperation, and gaze towards the opponent’s face. Across the population, we found that 33% of mSTS neurons distinguished between payoff conditions during the period when the tokens were presented (0-500ms from token onset), but only 14% of ACCg neurons did so (supplementary figure 3A). A small proportion of neurons in both areas encoded SPE estimated from the previous trial (10% in mSTS, 8% in ACCg).

We next focused analysis on the reward delivery epoch. Figure 4A shows an example mSTS neuron from a monkey playing a live opponent. This neuron fired more strongly for rewards received through cooperation than for equivalent rewards received for selfish actions (Figure 4ai). By contrast this neuron was much less sensitive to the amount of reward received (Figure 4aii). This neuron also fired strongly when the monkey predicted that the opponent had a low versus high probability of swerving on that trial (Pt). ((Figure 4aiii). We next asked whether mSTS neuron responses to cooperative reward might instead reflect perceptual social signals associated with looking at the opponent’s face. Overall, players tended not to look at their opponent’s face-space after cooperating, even after controlling for reward size (p<1 × 10^−40^). When we scrutinized only those trials on which the monkey did not look at his opponent’s face, this neuron still fired more strongly for cooperative rewards than selfish rewards (figure 4aiv).

We found similar neurons in ACCg when the monkey played a live opponent. An example neuron (Figure 4b) decreased its firing rate immediately after juice delivery for cooperation, but increased firing for non-cooperative juice rewards (Figure 4bi). This neuron also increased firing rate for ‘chicken’ rewards (Figure 4bii) but did not signal the monkey’s predictions of his opponent’s strategies (Figure 4biii). For those trials on which the monkey did not look at his opponent’s face, this neuron still differentiated cooperative and non-cooperative rewards (Figure 4biv).

Firing rates of between 29% and 47% of neurons in both brain areas were significantly modulated by the amount of realized reward in all three agency conditions during reward delivery (250-1250ms post juice delivery, between agency F=4.92, p=0.007), while 20-24% of them were similarly modulated in the post-decision period preceding juice delivery (Figure 4c). Remarkably, firing rates of 38% of mSTS neurons were modulated by cooperation, compared to only 20% in ACCg (2-way ANOVA, F=40.24, p<10^−7^) in the reward delivery period. Roughly 20% of neurons in both areas carried information about the opponent’s predicted strategy (Figure 4c). Activity of only a small percentage of neurons was modulated by gaze at the opponent (7-11% in ACC, 10-16% in mSTS).

We next explored in depth neurons that were selective for cooperation. We found that some of these neurons were excited by cooperation while others were suppressed (from figure 5a & 6a, categorized by the sign of the LM regression coefficient). When analyzed separately, subpopulations in mSTS distinguished the mechanism—cooperation or selfish choice—by which the same amount of juice was obtained (Figure 5a); the control trials where only one player played and could not crash, which had been held out from the LM analysis, showed distinct responses compared to the cooperative outcome but not the selfish outcome (yellow line, figure 5a & 6a). Similarly identified subpopulations in ACCg lacked these distinctive patterns of activation or inhibition in response to cooperation (figures 5b & 6b). Like neurons in mSTS, firing rates of ACCg neurons distinguished amount of juice received (Figure 4c), but showed no consistent differences between cooperative, selfish, chicken and control rewards (Figure 5b). Most importantly, population responses to cooperative rewards were no different than those for chicken outcomes, which were achieved by the same joystick movement. By contrast, mSTS neurons discriminated cooperative rewards from all others (selfish, chicken, and control; Figure 5a). This is especially noteworthy considering that joystick movement and subsequent car movements were perpendicular for the selfish and chicken outcomes, but cooperative and chicken outcomes were achieved by the same joystick movement and car translation. These findings strongly endorse the conclusion that mSTS cooperation signals are not mere reflections of sensory or motor task contingencies.

**Figure 5:**
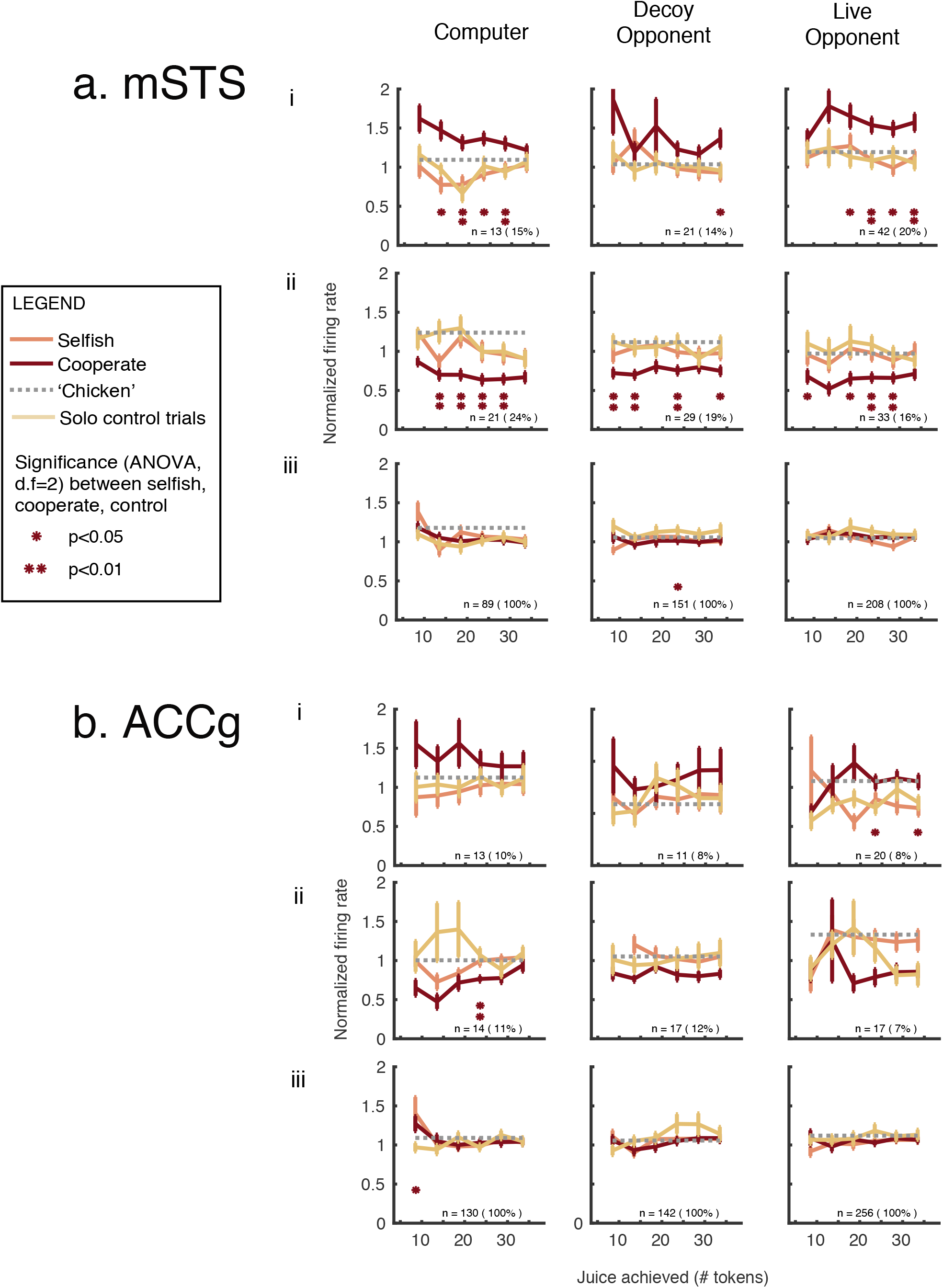
mSTS neurons selectively encode cooperation. A. Normalized mSTS population firing rate (250-1250ms after reward) segregated by excitation (top row) and suppression (middle) and all cells (third row). Error bars, +-SEM.
B. Normalized population firing rates in ACCg in same period. Conventions as in Figure 5A.

**Figure 6:**
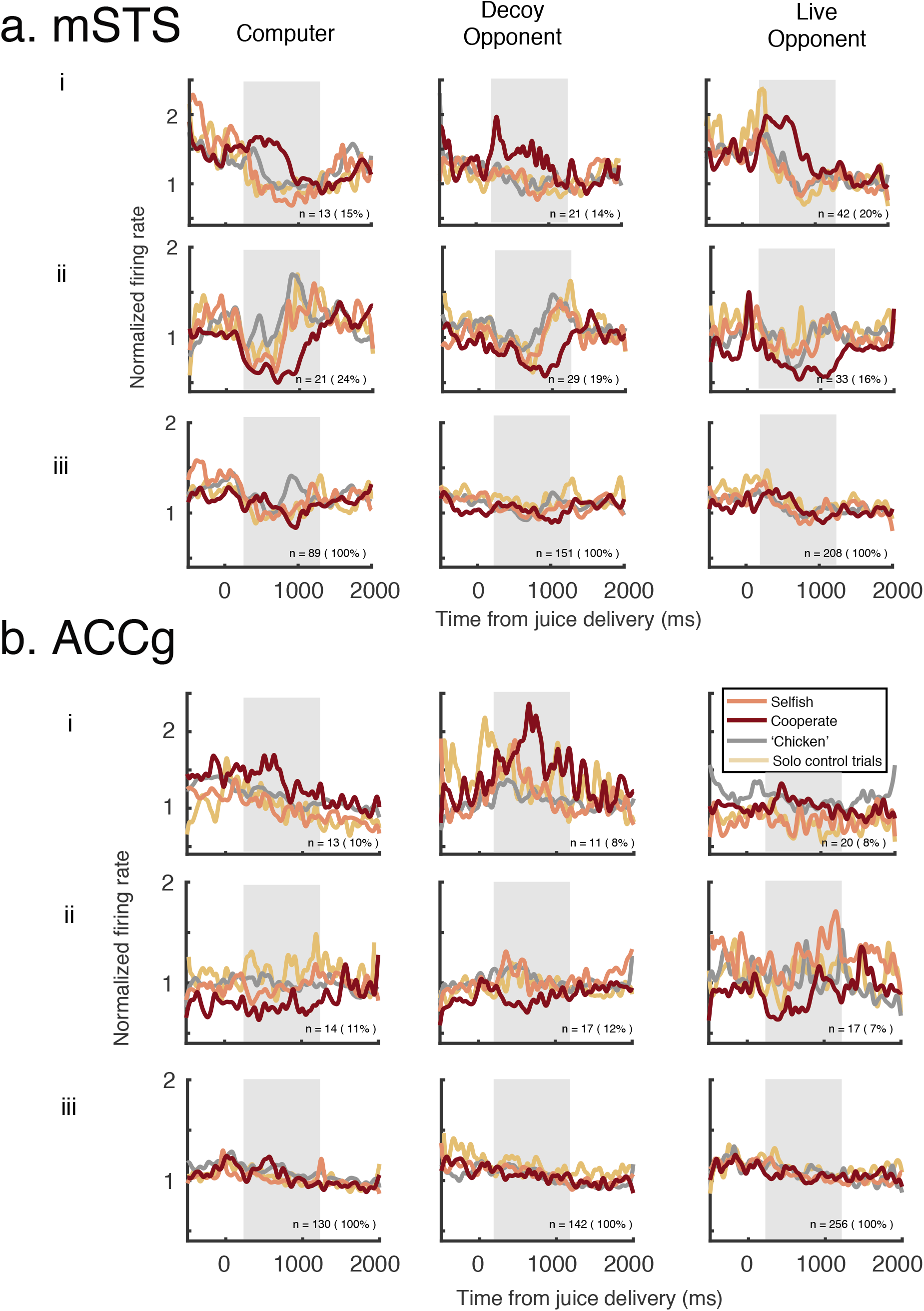
Neuronal cooperation signals vary over time. A. PSTHs for mSTS segregated by outcome. Rows show data for neurons enhanced (top), suppressed (middle), and all cells (bottom).
B. PSTHs for ACCg segregated by outcome. Conventions as in Figure 6A.

We focused the cooperative reward analyses on the epoch 250-1250ms after reward. Excitation and suppression by cooperation, however, varied over time. Early (250-750ms post-reward), 20-23% of mSTS neurons signaled cooperation by increasing firing rate, while 7-15% did so by decreasing firing rate. Later (750-1250ms post-reward), 25-29% of neurons decreased firing rate for cooperation while 7-15% of neurons increased firing rate (Supplementary Figure 4b). Overall, we found that 50-60% of mSTS neurons significantly encoded cooperation in one or both reward epochs (36-45% for a single epoch, 15-16% for both epochs), compared to a much smaller percentage of ACCg neurons (16-20% in a single epoch, 5-6% in both epochs; Supplementary Figure 4).

## Discussion

The evolutionary, economic, and biological origins of human cooperation remain hotly debated^26–28^. Both empathy and strategic reasoning contribute to cooperation in humans ^29^, supported by distinct but interacting brain networks. The evolutionary wellspring of human cooperation and the neuronal mechanisms that support it are not well-understood, in part due to the difficulty of eliciting strategic cooperation in animals, in which direct neural recordings can be made ^30^. To remedy this gap, we here show that monkeys understand and navigate a strategic game with payoffs that sometimes favor cooperation. Monkeys behaved as if they reasoned recursively about other individuals’ beliefs and desires in order to predict their choices and to guide their own actions, especially the decision to cooperate. They did not use simple strategies such as titfor-tat^22^ or win-stay-lose-shift ^23^ to play the game (Supplementary Figure 2b), nor could their behavior be explained by simple reinforcement learning. Monkeys paid close attention to payoffs available for both themselves and their opponents as well as intention signals indicating their opponent’s choice, and readily distinguished the agency of decoy and live players. These findings suggest monkeys implement a sophisticated model of their opponent in the game, and the recursive depth of this model varies with social status. Like humans ^31^, low status monkeys use skill and guile to interact strategically with higher status individuals ^32^, who are more likely to behave selfishly (Supplementary figure 1d).

Brain imaging studies in humans indicate that two interacting systems, one associated with empathy and social emotions and the other linked to mentalizing and social reasoning, support social interactions ^1,33,34^. Our findings show that neurons in putative primate homologs in both these systems, the ACCg and the mSTS, encode abstract information associated with strategic game play. Notably, non-perceptual social and strategic signals were stronger and more prevalent in mSTS than ACCg, and were sensitive to the agency of the opponent. By reverse inference ^35^, these findings endorse the importance of sophisticated reasoning in strategic interactions between monkeys revealed by our computational model.

Prior neurophysiological studies of STS revealed neurons that selectively respond to the sight of faces ^36,37^, facial expressions ^38^, and the direction of social gaze^36,39,40^. We found that roughly 20% of mSTS neurons were sensitive to looking at the face of an opponent, but these signals were weak (Figure 4C, D), suggesting either that perceptual social signals are dampened during strategic interactions or that different populations of mSTS neurons encode perceptual and abstract social information. In either case, we provide some of the first neurophysiological evidence for the representation of abstract, non-perceptual social information in primate mSTS, a finding that strongly endorses the hypothesis that this area is the homolog of human TPJ ^15^.

Though long thought to be uniquely human, the ability to strategically play mixed-motive games likely characterizes many social animals ^41,42^, particularly primates, that form differentiated relationships, including alliances and friendships, in order to navigate the complexities of group life ^43–45^. For long-lived, social primates like rhesus macaques, success depends on the deft deployment of cooperation and competition, which leverages individual identification ^46^, memory for previous interactions ^47^, investment of biological capital ^48^, learning ^32,49^, knowledge of others’ social relationships ^44,50^, and sensitivity to the quality of potential allies ^47^. Both prosocial behavior and cooperation in humans also depend on these factors, strongly suggesting the underlying mechanisms are conserved across anthropoid primates ^43^. Our findings confirm this prediction by demonstrating neurons in the primate social brain network—particularly mSTS—carry a wealth of non-perceptual strategic information, including payoffs, intention cues, and outcomes, and selectively signal rewards obtained by cooperation. Modulation of these signals by opponent agency further strengthens the similarity to human TPJ ^10,11^. Thus, large scale human societies, with all the complexity that attends cooperation and selfishness—whether in the boardroom or on the playground—arise from biological mechanisms that appear to have evolved early in the primate clade to support strategic social interactions.

## Methods

All experimental methods were approved by the Duke University Institutional Animal Care and Use Committee and were conducted in accordance with the Public Health Service *Guide to the Care and Use of Laboratory Animals*.

### Subjects

All procedures were approved by the Duke University Institutional Animal Care and Use Committee (protocol registry number: A295-14-12), and were conducted in compliance with the Public Health Service’s Guide for the Care and Use of Laboratory Animals.

Five male rhesus monkeys (8.6-13.2kg, 9-15 YO) were implanted with head-restraint prostheses (Crist) and neurophysiological recording cylinders (Crist) using standard sterile techniques as described previously (one of our papers here). Animals were initially anesthesized with ketamine hydrochloride and maintained with isofluorene (0.5-5% mg/kg). Enrofloxacin or other vet-prescribed broadspectrum antibiotics, and buprenorphrine for pain management were administered after surgical procedures. The animals were visually monitored continuously for at least 2 hours after surgery. The post-operative recovery period was 4 weeks, during which the animal was given free access to fluids and no training or testing was carried out. The recording chambers were cleaned at least 3x/week, treated with antibiotics and sealed with sterile caps. During testing, animals were given access to fluids amounting to at least 20mL/kg/day and supplemented with fruit and vegetables. Dominance relationships between pairs of monkeys were determined by controlled confrontation^51^.

### Behavioral task

Monkeys sat in primate chairs (Crist) facing each other at a distance of 30 inches with a 27 inch LCD screen placed horizontally between them (Figure 1a). Their heads were tilted slightly downward, at an approximate angle of 20°, allowing them to view both their opponent’s face and the screen.

Eye position and pupil diameter for one monkeys was sampled at 1kHz using an infrared eye tracker (SR Research Eyelink) mounted on the primate chair. At the start of each trial, the eyetracker sent timestamps to the experimental software (Matlab), which collated them with timestamps from the neurophysiological recording system (Plexon) and task events (PsychToolBox). The animals manipulated a joystick (60Hz) placed within the primate chair. The front of the primate chair, including the neck plate, was painted black to obscure the shoulders, hands, and joystick of both monkeys. The task was presented on a shared horizontal screen between the two animals. To initiate the task, the monkey whose eye position was being monitored had to fixate a central white dot on a black background (200ms). The fixation point was then extinguished, and two colored annuli (hereafter “cars”) appeared, one above and one below the extinguished fixation point. Each monkey controlled the car located closer to him, which was also cued by color (e.g., blue for M1 and red for M2; Figure 1b, Supplementary Figure 1a is an image with the task stimuli to scale). To continue, the joysticks for both animals had to be in the neutral position. After a variable delay <= 500ms, the coordination bar and four sets of tokens appeared, two for each player cued by color and position. The number of tokens was proportional to volume of juice available for achieving that option (0.02ml/token). Five hundred ms later, a moving dot kinematogram appeared within each car. Monkeys committed a choice by holding their joystick towards a specific token array for 500ms, at which point the white dots in the kinetogram changed to the player’s color (e.g., blue for M1 and red for M2); if they did not do so within 4 seconds, the last joystick direction was implemented as his choice. Monkeys were permitted to make multiple joystick movements freely as they deliberated their choice, which were immediately translated into the direction the dots moved within the car. After the choice period, the dots disappeared and the cars moved in the chosen direction. Juice rewards were delivered to the monkeys via a tube controlled by a solenoid valve. If the monkey achieved the token array straight ahead or the cooperative outcome, the solenoid opened twice, once for the smaller constant payoff and again for the second set of tokens in the same location.

The total number of tokens presented on the screen was always 41 for each player, divided between two locations, straight ahead and on the side of the screen. At each location, there was a small constant payoff of 3 tokens; the remaining 35 tokens were divided between the two locations in multiples of 5. The payoff for both animals was always symmetrical.

On 75% of trials, the larger reward was opposite the controlling monkey behind the opponent’s car; smaller rewards were to the side. To obtain the larger reward, M1 must go straight, but if M2 also did so straight their cars collided and neither received reward (*‘crash’*, Figure 1Fi). If M1 (M2) goes straight, receiving 28 tokens and M2 (M1) yields, receiving 3 tokens or the ‘chicken reward (Figure 1Fii & iii). On the remaining 25% of trials, the smaller reward was opposite the controlling monkey, behind the opponent’s car, and the larger rewards was to the side with all but 3 tokens behind the cooperation bar. To obtain the larger reward on these trials, both monkeys had to coordinate their movements and drive their cars to push the ‘coordination bar’(‘*cooperate’*) (Figure 1Fiv, Supplementary Figure 1). If only one monkey moved his car to the side and encountered the bar, it did not move and the monkey only received the 3 tokens in front of the bar (‘*chicken’* outcome).

On half of trials, the coherence of the moving dot kinematogram was randomized to obscure intention signals indicating the current directions in which the monkeys were holding their joysticks. Within session controls were trials on which only one monkey’s car and tokens were displayed. On control trials (10% total, randomly interleaved), monkeys should always choose straight regardless of the payout scheme since that will return at least 8 tokens while turning to the side will only yield 3 tokens. Control trials were excluded from the behavioral and neural analyses unless explicitly mentioned.

Seven monkeys were trained to play the task by first playing against a computer opponent that made straight/yield choices randomly. We varied dot motion coherence to ensure that the monkeys were attending to the projected future motion of the computer opponent’s car. Two animals did not reach criterion and were removed from further studies. The remaining 5 monkeys were deemed to have reached criterion when they were able to successfully avoid crashes 95% of the time when the intention signals were 100%. These monkeys also had thresholds, where their probability of crashing was 50%, that ranged between 15-25% dot motion coherence. Only two dot motion coherence levels were used in the final experiment: strong, 90% coherence; and weak, 0% coherence). All trials were randomly interleaved. The monkeys played against different opponents on consecutive days.

### Opponent agency conditions

‘*Live Opponent*’: Two monkeys were present in the experimental setup and both of them actively played the chicken game against each other (4 pairs, n= 75, 630 trials).

‘*Computer*’: One monkey was placed into the experimental setup opposite an empty primate chair. Joystick movements and choices from a randomly chosen prior live monkey behavior session were played back as an opponent to the current monkey, and any juice rewards obtained by the computer were delivered to the empty primate chair (4 players, n= 38, 938 trials).

‘*Decoy*’: Two monkeys were present in the experimental setup, but only one of the animals was the designated active player. The ‘decoy’ animal sat in the primate chair and drank the juice rewards delivered, but the joystick movements and choices from a prior live monkey behavior session were played back to the active player as the opponent (4 players, n= 49, 691 trials).

### Electrophysiological recordings

We acquired structural magnetic resonance images (3T, 1-mm slices) of each monkey’s brain. We made a mask consisting of a 3mm sphere around a seed at the fundus of the STS (X = 18.75, Y = −10.00, Z = −2.25) according to the Montreal Neurological Institute (MNI) atlas). This location in mid-STS was selected based on research indicating this region exhibits a functional connectivity profile most similar to the human TPJ (Mars et al., 2013). The mask was then converted into the individual monkey’s native-space structural scan to identify our target recording location (‘mSTS’) using FSL’s FMRI Expert Analysis Tool (FEAT) Version 6.0.0. For both mSTS and ACCg (Brodmann areas 24a and 24b), detailed localizations were made using Osirix (http://www.osirix-viewer.com) or Horos (https://horosproject.org) data viewer.

All single-unit recordings were made using single tungsten microelectrodes (FHC). In each recording session, a sterilized single electrode was secured onto the recording chamber (Crist Instrument) via an X-Y stage (Crist Instrument) and an adapter (Crist Instrument). The dura was penetrated using a sterilized guide tube (22 gauge, stainless steel, custom made), and the electrode was lowered through the guide tube via a hydraulic microdrive (Kopf Instruments). Signals were filtered and recorded using a 8-channel recording system (Plexon Inc). In addition to being guided by stereotaxic coordinates and MRI localization, each day we confirmed the recording site by listening to multiunit changes corresponding to gray and white matter transitions while lowering the electrode. For the STS recordings, we further verified the recording site by listening to multiunit activity that was visually responsive to a set of 200 images (consisting of human and non human primate faces, body parts, and objects). The neurons selected for recording in mSTS were within 150um of visually responsive cortex. Beyond that, neurons in both ACCg and mSTS were selected for recording based strictly on location, stability, and quality of isolation. A total of 528 ACCg and 448 mSTS neurons were recorded in 4 monkeys, in 3 agency conditions. Live opponents: 256 in ACCg, 208 in mSTS); decoy opponent (142 in ACCg 151 in mSTS); and no one (130 in ACCg, 89 in mSTS).

### Analysis of behavioral data

All behavioral data, including joystick movements and eye-tracking data, was collected and analyzed with custom code on MATLAB.

For analyzing event outcomes over time, trials from all sessions were collated into 5-trial bins for the same agency condition (blue, live opponent; green, decoy; grey, computer, figure 2A). To look at the event outcomes over payout conditions (difference in the number of tokens available straight ahead, Vstr, and cooperate, Vcoop), the trials were sorted into high signal trials where players’ joystick movements are indicated by moving dots in the cars; and low signal trials in which dots moved randomly.

For analyzing eye position signals, we drew boxes around the areas of interest (supplementary figure 1a) and quantified the instances in which eye position fell into those areas. The face region of the recipient was determined empirically prior to the experiments and defined as the area between the neck plate and the top bar, and the side panels of the primate chair. We used a large window to capture gaze shifts that were brief in duration and large in magnitude and often directed at varying depths (e.g., eyes, mouth). In the empty chair condition, this would be the space where the opponent’s face would have been^52^.

Eye positions were plotted in 1ms bins, and shown on the figures with standard errors of the mean calculated between behavioral sessions. The trials were also sorted into high and low signal trials, and the difference in looking behavior between the conditions indicate a bias.

Statistical tests were conducted as two-tailed ANOVAs with multiple comparisons (Tukey’s HSD test) unless otherwise specified. All figures are shown with standard errors of the mean unless otherwise noted.

The hybrid reinforcement learning-strategic learning model was computed using Stan^53^ via the MATLAB interface. The choice behavior from each pair of opponents and each agency condition were fit separately. The model predicts the choices of one animal conditioned on their opponent’s choices, so the data from each monkey pair was fit twice; once with a given monkey being the agent whose choices we were predicting and a second time with that same monkey being the opponent. All model comparisons were performed using the Akaike Information Criterion (AIC)^54^. To compare goodness-of-fit across different subsets of the choice data, we use the log-likelihood per trial (Figure 3B, Supplementary figure 2), as the absolute log-likelihoods are strongly influenced by the number of trials in the data set.

### Model specification and fitting

#### a) Hybrid reinforcement learning and strategic learning model

We model each animals’ choice behavior using a model that combines a simple reinforcement learning (RL) system with an expected value model. The RL system takes into account only which action, straight or yield, the animal has taken and what reward they receive, while the expected value system prospectively takes into account which potential reward outcomes are available on each trial (as indicated by the token symbols on the game board), and chooses according to how likely each outcome is.

The probability of the animal yielding on trial *t* is determined by the difference in utility between the yield and straight choices, *U*_*t*_, according to the equations

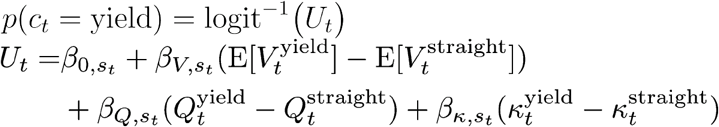

where *c*_*t*_ ∈ {yield, straight} is the animal’s choice on trial *t*. The utility difference is a linear combination of the output of three valuation sources, denoted by the *Q, κ* and E[*V*_*t*_] values respectively, each weighted by a temperature parameter. We will discuss each of these sources of valuation in turn. Note that the temperature parameter for each system differs depending on the signal strength *s* used on the trial *t*, which can be either high or low, for a total of six temperature parameters.

The *Q* values for yield and straight are learned through a simple RL system using reward prediction error update equations,

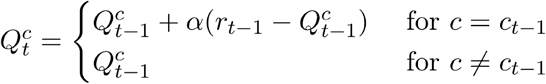

Here the *Q* value of the previously chosen choice is incremented according to the learning rate *α* towards the reward received. At the beginning of a session both *Q* values are initialized to the value *Q*_0_, which is fit as a free parameter bounded between zero and the largest possible payoff on any trial.

Second, the *κ* values capture autocorrelations in the animal’s choices, such as a tendency to either repeat or avoid the action that has been taken recently. Similar to the *Q* values, the *κ* values are updated each trial using a Rescorla-Wagner rule,

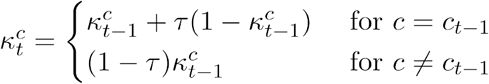

where the parameter *τ* ∈ [0, 1] determines how rapidly the influence of past choices decays. Each *κ* value is initialized to zero at the beginning of a session.

Finally, the expected reward values 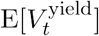 and 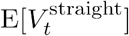 are estimated by the animal on each trial based on the potential reward values indicated on the game screen as well as the animal’s beliefs about his opponent’s strategy. On each trial the animal will obtain one of four possible reward values, 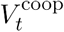 if both the animal and his opponent yield, 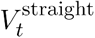 if the animal chooses straight while his opponent yields, *V*^safe^ if the animal swerves while his opponent goes straight, or *V*^crash^ if both the animal and their opponent go straight. Note that the safe and crash values do not change from trial to trial and are fixed at three tokens and zero tokens, respectively. Accordingly, the animal calculates the expected values of the actions straight and yield given his belief regarding how likely his opponent is to yield. This likelihood is denoted by 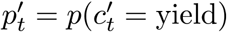 where 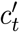 is the opponent’s choice on trial *t*. The formulae for the expected values are given by

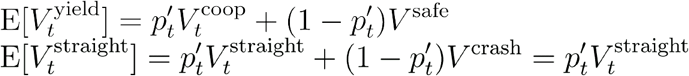

This prompts the question of how the animal obtains his beliefs regarding his opponent’s strategy – that is, where does *p*′ come from? In our model, the animal’s representation of his opponent’s strategy takes the form of a logistic regression which maps the characteristics of each trial onto a probability of yielding. The animal learns this logistic regression using online-updating as he observers his opponent’s actions. Formally, this is given by

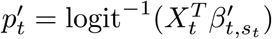

where 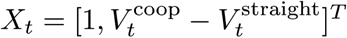 is a vector of regressors consisting of an intercept and the difference between the cooperative and straight reward values, and 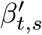 is a vector of regression coefficients. Note that the animal’s belief regarding his opponent’s strategy differs between the high and low signal conditions, as 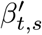 differs depending on the signal strength *s*.

The animal updates his beliefs about his opponent’s strategy on each trial using stochastic gradient descent updates given by

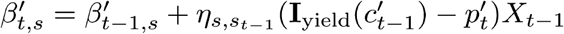

The logic of this update is very similar to that of reward prediction error (RPE) updating used in RL models. In an RL model, the predicted reward value *Q* is updated such that it will be closer to the reward received on the previous trial. Analogously, here the regression coefficients are updated such that the prediction of the logistic regression *p*′ will be closer to the outcome observed on the previous trial. The size of the step taken towards the previously observed value is governed by a learning rate, here denoted *η*.

As in an RL model, the critical quantity for trial-by-trial learning in our strategic learning model is the error term that captures how predictions differed from the true outcome. Here this error is the term 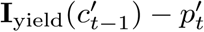, where **I**_yield_ (*c*′) is the indicator function that returns one if the opponent’s chose yield and zero if they chose straight. We refer to this quantity as the strategic prediction error (SPE) in analogy to the RPE of RL systems.

Intuitively, beliefs about the opponent’s strategy on low signal trials may be less affected by trials from the high signal condition, and vice versa. Therefore, the regression coefficients for the high and low signal conditions, 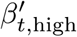 and 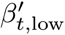, are updated differentially depending on which signal condition of the previous trial. Specifically, each set of regression coefficients has different learning rates depending on whether the trial that is being learned from was high or low signal condition, such that 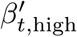 is updated using *η*_high,high_ if the previous trial was high signal condition, and using *η*_high,low_ otherwise. The same is true for 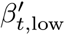, for a total of four different learning rates.

Beliefs about the opponent’s strategy at the beginning of a session are determined by the initial values 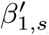, which are fit as free parameters.

#### b) Submodel comparisons

In order to determine the level of sophistication of each animal’s choice behavior, we compare a number of submodels of the model presented above, as well as the full model. The full model assumes that the player ascribes intention (or theory of mind) to the opponent and represents his opponent’s strategy in the form of a logistic regression and updates it online via the strategic prediction error (SPE). Each submodel is equivalent to the full model with a subset of those features turned off, which we accomplish by fixing certain parameters at zero. We describe the submodels in order of (approximately) increasing sophistication, and in the same order that the submodels are shown, from left to right, in the x-axes of figure 3B.

1. The least sophisticated submodel is a naive RL model in which all *β* parameters other than *β*_*Q*_ and *β*_*κ*_ are fixed at zero. This model estimates the values of the actions swerve and straight based only on reward history and does not incorporate the visual information presented on each trial about the payoffs available.
2. The next submodel is a logistic regression on the payoffs available on the current trial. This model does not use RL or SPE and instead chooses based only on the visual presented about the payoffs on each trial. *β*_*Q*_ all learning rates *η* are fixed at zero. Also, the second elements of the parameter vectors 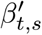 are fixed at zero, which leads to beliefs about the opponent’s strategy being invariant to payoff condition; i.e. the animal does not consider that their opponent has their own intentionality and cares about the payoff condition.
3. A combined logistic-RL model, equivalent to the second model described above with *β*_*Q*_ not fixed at zero.
4. An model that incorporates SPE learning, but without representing the opponent’s intentionality. This is equivalent to the full model with the second elements of the parameter vectors 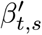 fixed at zero.
5. A ‘static’ ToM model where the opponent is assumed to have intentionality and cares about obtaining higher payoffs, but there is no SPE learning and so beliefs about the opponent’s strategy do not adjust over time. This is equivalent to the full model with all learning rates *η* fixed at zero.

Using the expected utility calculations, payoffs and opponent’s predicted behavior, we were able to compute and predict players’ behavior as well as his prediction of his opponent’s behavior (Figure 3C, hybrid RL-logit and ToM for an example pair).

### Analysis of electrophysiology data

Single-unit activities were isolated using a combination of principle component analysis (PCA), the Template Matching algorithm, and hand-sorting in Offline Sorter (Plexon Inc). All subsequent data analyses were accomplished with custom MATLAB scripts. The peristimulus time histograms (PSTHs) shown are rendered in 1ms steps with Gaussian smoothing of 10ms on both sides. For population PSTHs, firing rates were normalized to the pre-fixation firing rate (200ms time window immediately before the onset of the fixation cue). Using different time windows and an alternative normalizing methods of a) normalizing to whole trial firing rates, and b) z-scoring of firing rates to the whole trial did not significantly change any main results reported. Statistical tests were conducted as two-tailed ANOVAs with multiple comparisons (Tukey’s HSD test) unless otherwise specified.

Epoch-based analysis were conducted for 3 distinct time windows: payoff presentation (0-500ms after the onset of the tokens on the screen), post-decision/cars move (0-500ms after the end of the 4s decision period and start of car movement), and juice delivery (250-1250ms after the juice is delivered).

The responses (neuronal firing rate in the epoch of interest as described above) from non-control trials were fit with linear models (LM). All continuous variables (including neural responses) were z-scored by the trials that made up each neuron’s data structure. The models were individually fit to each neuron and the and ANOVAs were used to classify the responses. The variables used in each model are as follows: For payoff presentation, the differences between the straight and cooperative token amount (Vdiff), the predicted strategy of the opponent for the current trial (Pt), and the strategy prediction error for the trial immediately prior (SPE1), all of which are continuous variables. For the post-decision/cars move and juice realization epoch, we include categorical variables as follows: cooperate, signal strength (indicating availability of explicit information about intentions), and gaze (considered ‘1’ if the animal makes a fixation for 150ms or longer within the defined ‘face’ boundaries of his opponent during the 1.5s window from juice delivery), and the continuous variable reward amount, which is the number of tokens the player received in juice. The opponent’s predicted strategy (Pt), a continuous variable, is orthogonalized against ‘cooperate’ and signal strength to avoid collinearity of the model, and is shown on its own as well as an interaction term with cooperate.

The terms used to categorized outcomes in figures 5 and 6 are *‘cooperate’*, where both players moved pushed the cooperation bar and received the Vcoop payout; *‘selfish’*, where one player goes straight (the *selfish* player), and the other deviated (‘*chicken’*); and the controls, where only one player’s avatar was present.

## Supporting information

all supplement

## Data availability

The data that support the findings of this study are available from the corresponding author upon reasonable request.

## Code availability

The custom analysis code for this study are available from the corresponding author upon reasonable request.

## Acknowledgements

We thank the DLAR staff at Duke University for providing excellent animal care. This work was supported by: R01MH095894, R01MH108627, R37MH109728, and a grant from the Simons Foundation (SFARI 304935, MLP).

## Author contributions

WSO conceived and carried out the experiments, and analyzed the data. SM-K designed the behavioral model. WSO and MLP wrote the manuscript with input from SM-K.

## Author information

The authors declare no competing financial interests.

