## Supplementary material for "Neuronal Mechanisms of Strategic Cooperation": all supplement

Extended data

Figure legends

SI Figure 1

A) Heatmap of fixation points for one session. The player looked at his opponent's car, his own car, the number of tokens in each location, and his opponent's face.
B) Top: probability gaze direction during various task phases from: the token onset, dot onset (decision period), cars move (review of outcome), and juice (realization of outcome)
C) Probability of gaze direction was directed towards the opponent's face synchronized on intention cue onset in live (blue), decoy (green), or computer (black) agency conditions. Second row: probability gaze was directed towards the opponent's car. Third row: probability was directed towards the tokens straight ahead. Bottom row: Probability gaze was directed towards the cooperative tokens.
D) Probability of gaze direction towards the opponent's face following juice for cooperative (red), selfish (orange), and chicken (grey) reward outcomes when playing a live opponent.
E) Scatterplot of frequency of yielding vs. frequency of choosing straight for 4 monkey pairs segregated by social dominance. For a given pair of players, we show the number of times they choose to yield (y-axis) or to play selfishly by going straight (x-axis). The points that lie below the diagonal indicate that those are the players that picked to go straight more than yield, while the points that lie above indicate the players that yielded more. We find that this corresponded to the relative dominance between the two players, The marker colors indicate player identity, while the color of the marker outlines (green, red, yellow, blue) indicate the players that are playing against each other. Note that the mid-ranking players (purple and brown) have points that fall on both sides of the diagonal, suggesting that their play strategy is not most consistent with their identity, but consistent with the dominance hierarchy depending on who the opponent is.

### Supp Figure 1

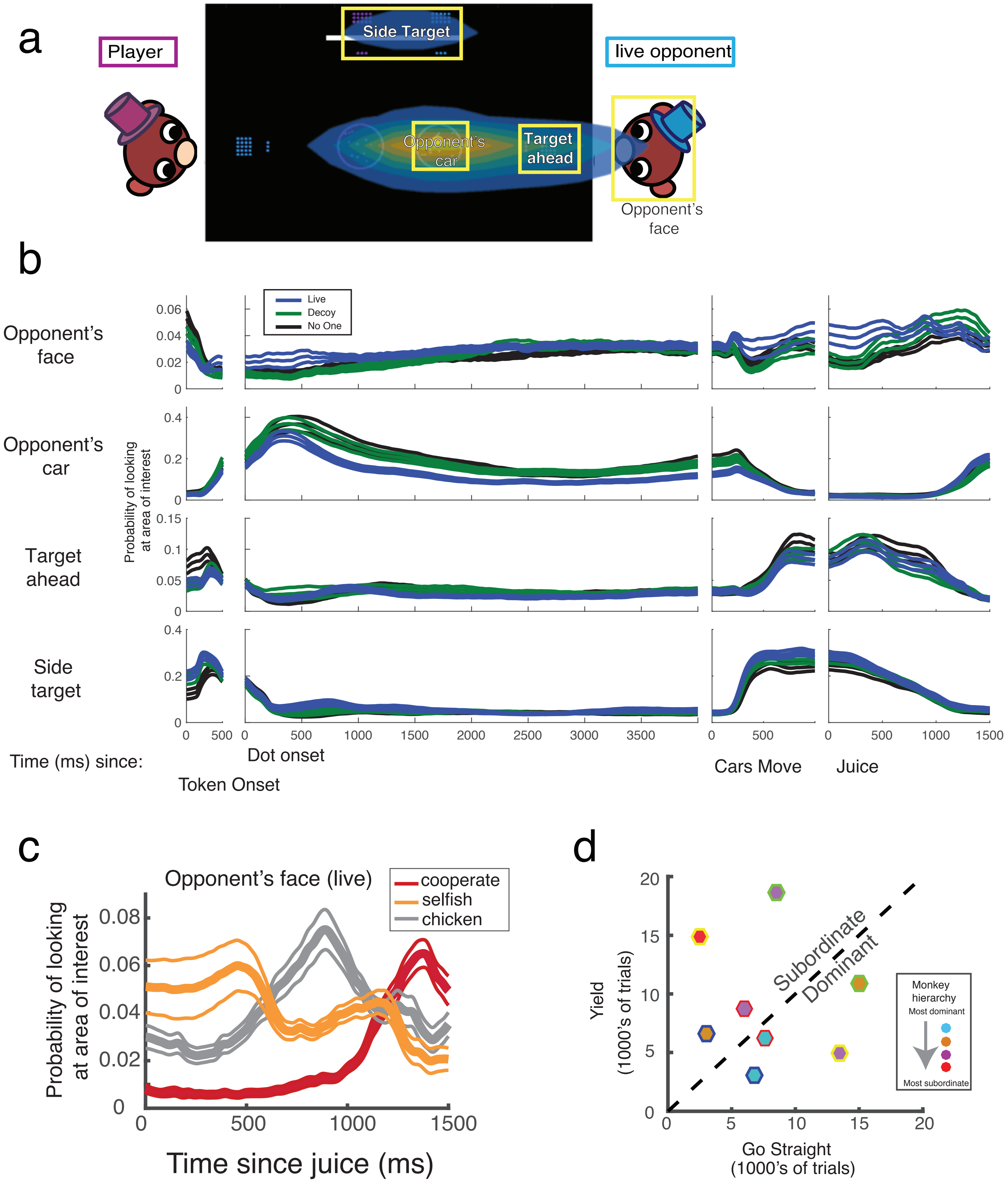

SI Figure 2

A) Average (grey bars) and individual monkey (colored points) log-likelihood ratio per trial (LLR) and Akaike weights(Wagenmakers & Farrell 2004) for each model (light pink bars). Models: 1. RL-logit in which payout conditions (Vt) were not included; 2. RL-logit in which reward prediction error (Q) was not included; 3. RL-logit model; 4. RL-logit model with strategic prediction error (SPE); 5. RL-logit model with intentionality/recursion (k1); 6. RL-logit model with intentionality/recursion (k1) and SPE.

B) Top row: probability players used a tit-for-tat strategy. First column shows predicted choices for the current trial if opponent chose to yield (black) or go straight (red) on previous trial. This strategy predicts the player will repeat the opponent's previous choice independent of payouts (reward difference, x-axis). Middle column assumes players only base choices on previous trial. Third column assumes the players treat different payout conditions independently and choose based on the last trial with the same payouts (reward difference).

Bottom row: probability players used a win-stay-lose-switch strategy. First column shows probability player repeated choice made on previous trial if he won (black) or lost (red). This strategy predicts the player repeats prior choices that lead to wins and switches from choices that lead to losses independent of payout conditions (reward difference, x-axis). We define 'winning' as opponent yielding. Middle column assumes the players only base choices on previous trial outcomes. Third column assumes the players treat different payout conditions independently and based on the last trial with the same payouts (reward difference).

### Supp Figure 2

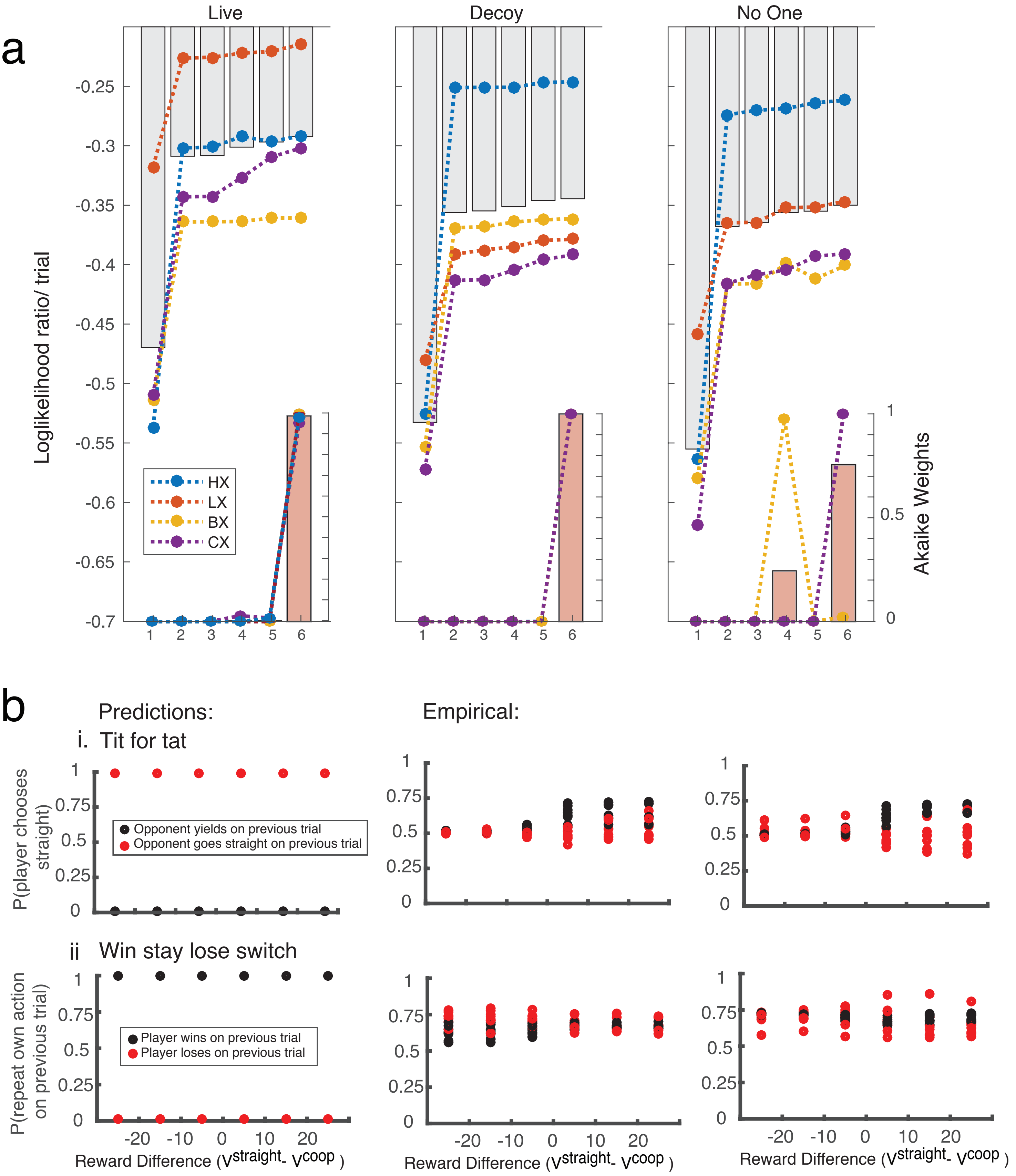

SI Figure 3

A) PSTHs for 2 example neurons and 2 example mSTS neurons in monkeys playing against a live opponent aligned to onset of moving dots (left column) and juice delivery (right column). Token onset occurs 500ms before the dots onset, cars move 4000ms later and juice delivery occurs 900 +/-100ms after cars move. Events marked by dotted lines. Top two rows depict neurons sensitive to payouts (light blue: Vstraight > Vcooperate; dark blue: Vcooperate > Vstraight); bottom two rows depict neurons sensitive to intention signals (dark red, low signal; light red, high signal).

B) Mean coefficients and percentage of significant cells returned by linear models (LR) of firing rates in the 500ms period following token presentation with 3 independent variables: payoff difference between options , Vdiff; probability opponent yields on current trial, Pt; and strategic prediction error from preceding trial, SPE1. Models were fit to data from each neuron separately and the results aggregated by recording location (mSTS, green; ACCg, red) and agency condition (color intensity: light (computer), medium (decoy), dark (live opponent)).

### Supp Figure 3

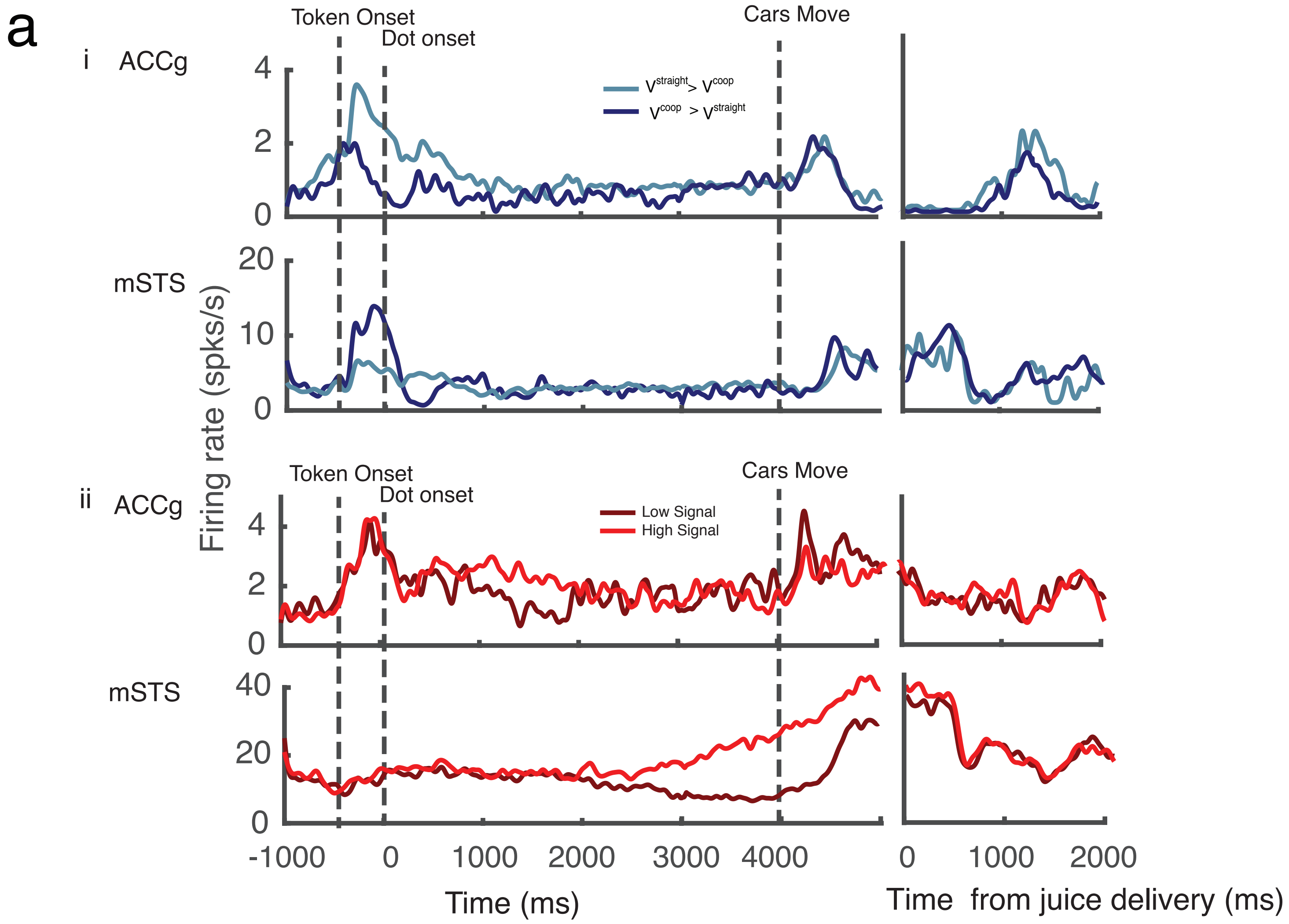


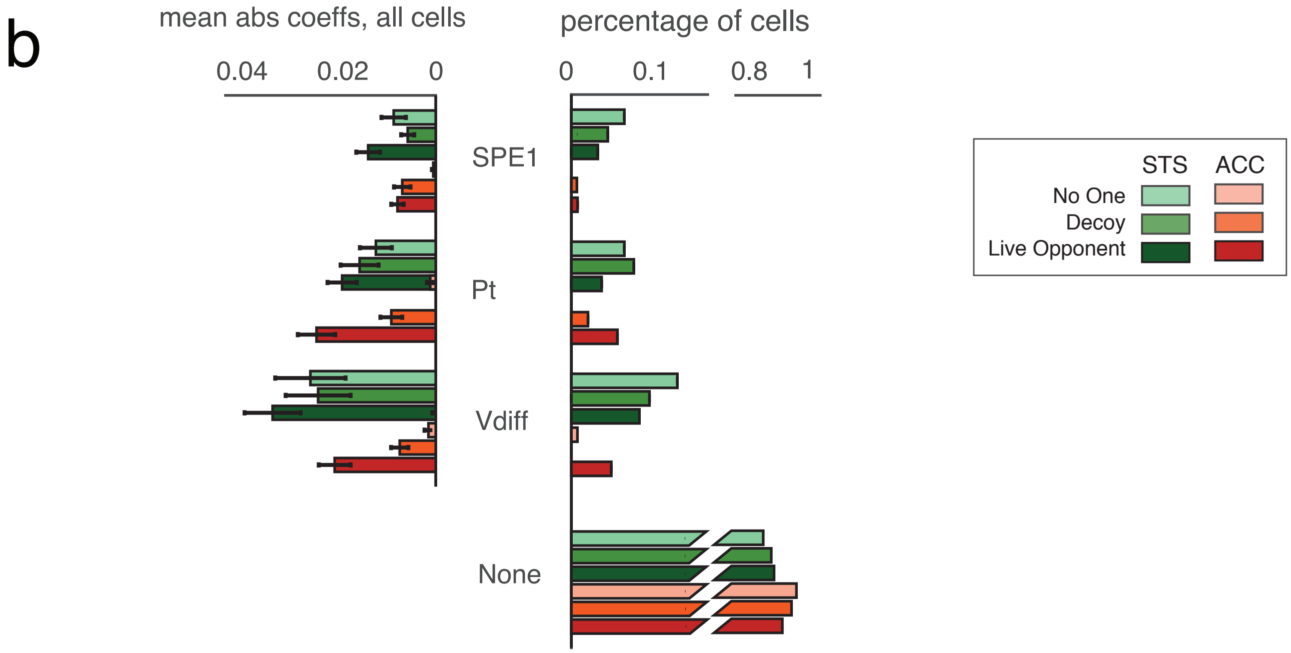

SI Figure 4

A) LM beta values for ‘cooperation’ in early (250-750ms) and late (750-1250ms) juice delivery period, as in Figure 4. Neurons not significantly modulated by cooperation in either epoch (grey circles), neurons significantly modulated in one epoch (black circles), and neurons significantly modulated in both epochs (red circles) plotted for mSTS (top panel) and ACCg (lower panel).
B) Percentage of neurons significantly enhanced or suppressed by cooperation in early and late reward delivery periods in ACCg and mSTS.

### Supp Figure 4

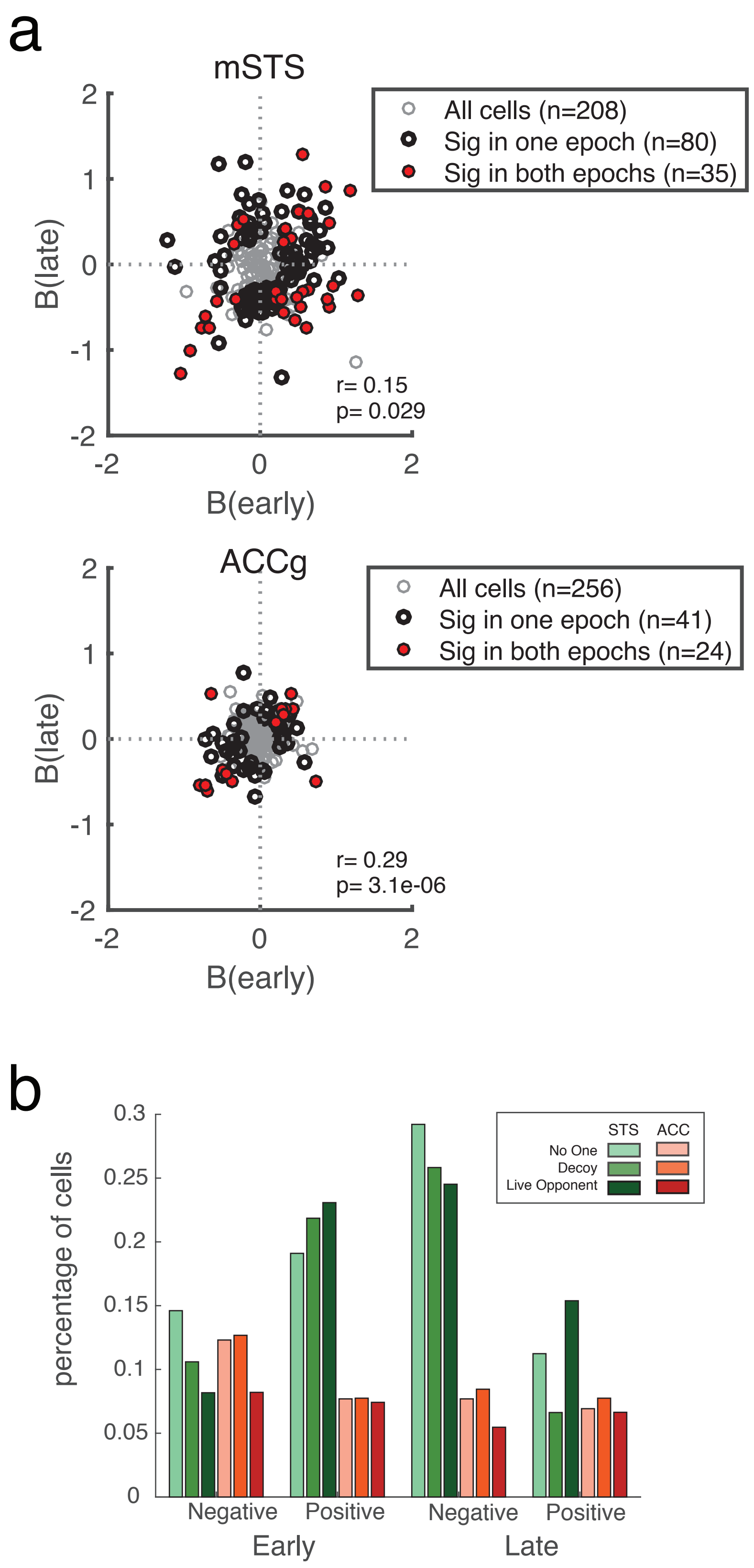

SI Video (Task)

Two players (Player 1,purple; Player 2, blue) sat opposite each other across a shared horizontal screen. The Player 1 controlled the purple annulus ("car") with a joystick. He could choose to go straight ahead behind player 2's (blue) car, or 'yield' and move to his left. In the first trial, he swerves and receives only 3 tokens as player 2 chose straight; in the second trial both swerved/yielded and "pushed" the white coordination bar together, displacing it and allowing access to the tokens behind it. On the 4th trial both players chose straight and collided, neither receiving reward. In the first 3 trials, the joystick movements are indicated by moving dots in the car (high signal trials); in the 4th trial the moving dots moved randomly (low signal trials). For more details see methods. Each token was worth 20ul of juice.

Supplementary Table 1

Comparisons between the classifications of neuron selectivities between areas (ACCg, mSTS), and the opponent conditions (live, decoy, computer). F-statistics and the corresponding p-values from 2-way ANOVAs are shown below

a) 0-500ms after car movement

| Cars Move | Proportion of cells | | Model fit parameter | | |
|---|---|---|---|---|---|
| | F-statistic | p-value | F-statistic | p-value | |
| **Cooperate** | 4.0829 | 0.0436 | 27.35 | 2.08E-07 | **Brain area** |
| | 1.0272 | 0.3584 | 2.2 | 0.1111 | **Opponent** |
| **Opponent's predicted strategy, Pt** | 5.19 | 0.023 | 20.33 | 7.34E-06 | **Brain area** |
| | 0.15 | 0.8573 | 0.29 | 0.7471 | **Opponent** |
| **Reward amount** | 0.86 | 0.3552 | 4.29 | 0.0387 | **Brain area** |
| | 1.47 | 0.2296 | 5.74 | 0.0033 | **Opponent** |
| **Signal** | 13.59 | 2.39E-04 | 6.52 | 0.0108 | **Brain area** |
| | 3.19 | 0.0417 | 4.35 | 0.0132 | **Opponent** |
| **Gaze** | 0.02 | 0.8808 | 11.95 | 5.71E-04 | **Brain area** |
| | 0.08 | 0.9214 | 1.61 | 0.1998 | **Opponent** |

b) 250-1250ms from juice delivery

| Juice delivery | Proportion of cells | | Model fit parameter | | |
|---|---|---|---|---|---|
| | F-statistic | p-value | F-statistic | p-value | |
| **Cooperate** | 39.03 | 6.23E-10 | 92.08 | 7.11E-21 | **Brain area** |
| | 0.86 | 0.4219 | 5.25 | 0.0054 | **Opponent** |
| **Opponent's predicted strategy, Pt** | 9.48 | 0.0021 | 9.21 | 0.00225 | **Brain area** |
| | 4.1 | 0.0169 | 1.49 | 0.2261 | **Opponent** |
| **Reward amount** | 3.02 | 0.0826 | 38.19 | 9.50E-10 | **Brain area** |
| | 4.46 | 0.0118 | 0.62 | 0.5361 | **Opponent** |
| **Signal** | 0.92 | 0.34 | 2.76 | 0.0968 | **Brain area** |
| | 0.01 | 0.99 | 0.13 | 0.8757 | **Opponent** |
| **Gaze** | 3.45 | 0.0637 | 11.81 | 6.00E-04 | **Brain area** |
| | 4.35 | 0.0131 | 0.49 | 0.0834 | **Opponent** |
